# *Cardinium* symbiosis as a potential confounder of mtDNA based phylogeographic inference in *Culicoides imicola* (Diptera: Ceratopogonidae), a vector of veterinary viruses

**DOI:** 10.1101/2020.09.20.305342

**Authors:** Jack Pilgrim, Stefanos Siozios, Matthew Baylis, Gert Venter, Claire Garros, Gregory D. D. Hurst

**Author notes:** Address correspondence to Jack Pilgrim.

## Abstract

*Culicoides imicola* (Diptera: Ceratopogonidae) is an important Afrotropical and Palearctic vector of disease, transmitting viruses of animal health and economic significance. The apparent incursions of *C. imicola* into mainland Europe via wind-movement events has made it important to trace this species to better predict new areas of arbovirus outbreaks. A widely used method for tracking dispersal patterns of *C. imicola* employs a phylogeographic approach anchored on the mtDNA marker *COI* (cytochrome c oxidase subunit I). However, a problem with this approach is that maternally-inherited symbiotic bacteria can alter the frequency of *COI* mitochondrial haplotypes (mitotypes), masking the true patterns of movement and gene flow. In this study, we investigate possible associations of the symbiont *Cardinium* with *C. imicola* mitotype distribution. Haplotype network analysis indicates the concordance of specific mitotypes with *Cardinium* infection status in *C. imicola* populations from the Mediterranean basin and South Africa. This observation urges caution on the single usage of the *COI* marker to determine population structure and movement in *C. imicola*, and instead suggests the complementary utilisation of additional molecular markers (e.g. microsatellites and nuclear markers).

## Introduction

*Culicoides* spp. (Diptera: Ceratopogonidae) are a group of tiny biting midges responsible for the spread of several pathogens to animals, including African horse sickness, Schmallenberg and bluetongue viruses. An important vector species, *Culicoides imicola*, was initially considered to be limited to Afrotropical regions but recent genetic analysis suggests this species has been present in the Mediterranean basin since the Late Pleistocene or Early Holocene (1). The movement of *C. imicola* at the northern fringes of the Mediterranean basin is likely driven by wind-events which have led to recurrent incursions of the species into mainland Europe (2). Thus, the tracking of *C. imicola* through population genetics and climatic modelling is important in establishing where potential future outbreaks of midge-borne viruses may occur.

Mitochondrial DNA (mtDNA) markers have previously been used in phylogeographic studies to detect biodiversity and dispersal events of *C. imicola* populations (1–3). However, mtDNA based inferences can be confounded by the linkage disequilibrium (co-inheritance) of symbionts and host mtDNA. As symbionts enter a naïve population, selective sweeps of the bacteria can occur and with it the homogenisation of linked mtDNA haplotypes, leading to the apparent absence of biodiversity (4). Conversely, variation in symbiont presence over space can lead to a perceived genetic population structure when none exists. Subsequently, inferences from these studies can only be interpreted in the context of any history of infection with symbiotic bacteria.

In this study, we examine the relationship between mtDNA in *C. imicola* and the heritable symbiont *Candidatus* Cardinium hertigii (Bacteroidetes). *Cardinium* is present in several biting midge species, including *C. imicola*, although *Cardinium*’s significance in *Culicoides* biology is unknown (5–7). As *Cardinium* infection could have implications for inferring movement of *C. imicola* into naïve geographic areas through mtDNA marker analysis, we analysed associations between mitochondrial haplotypes (mitotypes) and *Cardinium* infection in this important vector species.

## Materials and methods

106 *C. imicola* individuals were collected between 2015-2016 using light traps from two countries: France (location: Corsica) and South Africa (location: Onderstpoort). Amplification of the host *COI* gene (8) and the *Cardinium* Gyrase B (*GyrB*) gene (7) was undertaken by conventional PCR assay. DNA extracts which passed quality control through successful *COI* amplification but were negative for *GyrB* were then screened with a more sensitive nested PCR assay (cycling conditions and primer sequence information in Table S1). PCR assays consisted of a total of 15 μL per well, comprising of 7.5 μL GoTaq^®^ Hot Start Polymerase (Promega), 5.1 μL nuclease free water, 0.45 μL forward and reverse primers (concentration 10 pmol/μL) and 1.5 μL DNA template. PCR products were separated on 1% agarose gels stained with Midori Green Nucleic Acid Staining Solution (Nippon Genetics Europe). *COI* amplicons identified by gel electrophoresis were purified enzymatically (ExoSAP) before being sent for sequencing through the Sanger method (GATC Biotech AG, Konstanz, Germany). Chromatograms were then visually assessed for correct base calls. Accession numbers for *COI* sequences generated in this study are LR877434-LR877461.

The mtDNA structure of *C. imicola* was then analysed in relation to *Cardinium* infection status. To this end, haplotype networks were constructed, with analysis and visualisation undertaken using the TCS haplotype network algorithm generated in PopART v1 (9). The networks included *C. imicola* mitotypes from this study (South Africa and Corsica), as well as mitotypes detected from the Iberian peninsula and Israel (Dallas *et al*., 2003; Portugal n=12, Israel n=29; Accession numbers: AF078098–AF078100, AF080531-AF080535, AJ549393–AJ549426). The Iberian population is known to be free from *Cardinium*, whereas that of Israel is known to carry *Cardinium* (5,6).

## Results/Discussion

To confirm individuals in this study were not cryptic species of *C. imicola* missed by morphological identification, a distance estimation of mtDNA barcodes was assessed giving a minimum identity of >98%, consistent with all individuals belonging to the same species. *Cardinium* infection was observed in the South African population (10/33 individuals) but not in Corsican populations (0/73 individuals) (Table 1). The additional nested screening revealed no evidence of low-titre infections. The absence of *Cardinium* in *C. imicola* populations from Corsica recapitulated a recent study from Spain (6), suggesting a lack of infection in populations from the Western Mediterranean basin. However, the presence of *Cardinium* in Israel (5) indicates infection heterogeneity exists between the Eastern and Western Mediterranean basins (EMB and WMB).

**Table 1.**
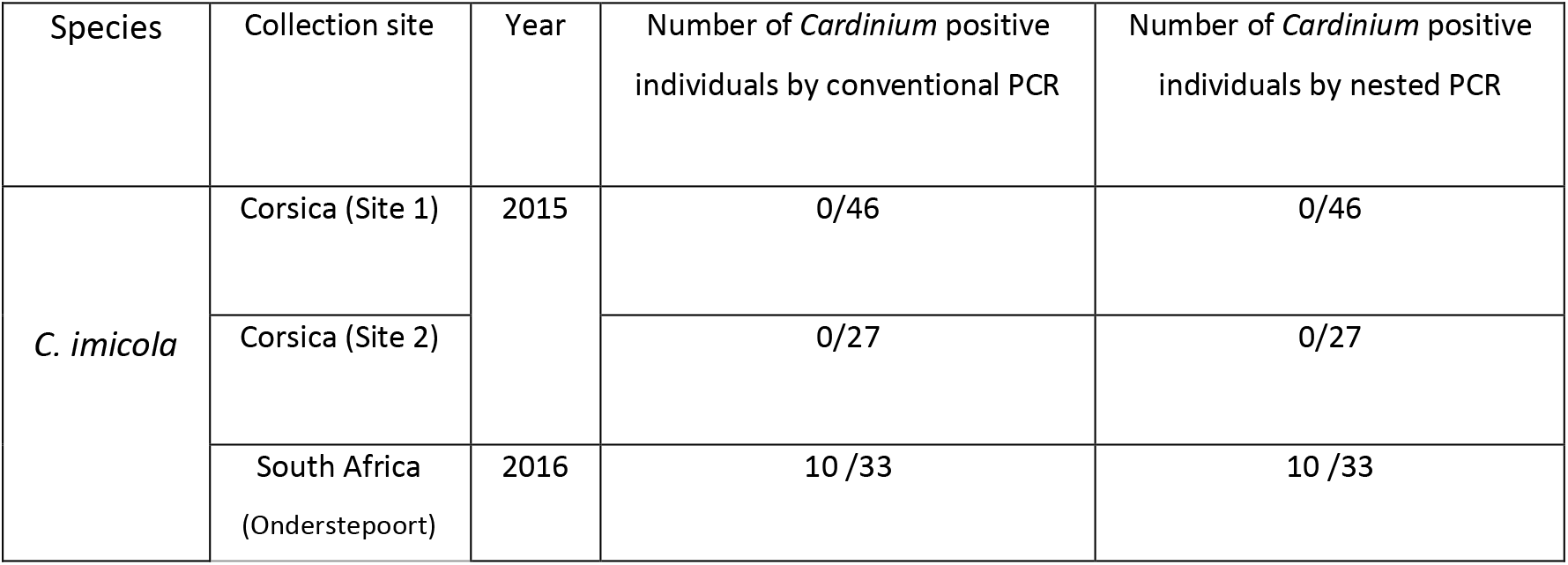
*Cardinium* screening results using conventional and nested *GyrB* PCR assays.

To investigate associations between symbiont and mitotype, mtDNA haplotypes and the *Cardinium* infection status of *C. imicola* individuals within and between populations were compared. A *C. imicola* mitotype network (Figure 1A) of the single South African population, containing both *Cardinium-infected* and uninfected individuals showed the presence of seven mtDNA haplotypes split into two broad mitotype groups separated by three SNPs. *Cardinium* infection status and these general mitotype groupings were associated, with all *Cardinium-infected* specimens falling in one group (Fisher’s exact test, *P*<0.001). As both symbiont and mitochondria are maternally inherited together, a selective sweep of *Cardinium* is likely to have led to this concordance of mitotype and *Cardinium*. Null models of symbiont-mtDNA dynamics predict three phases. First, the symbiont sweep, during which the symbiont associated mitotype increases in frequency, and the infected and uninfected individuals have different associated mitotypes. This is followed by the mitotype of infected individuals becoming increasingly represented in uninfected individuals where the symbiont fails to transmit from mother to offspring. In the final phase, mutation re-establishes diversity in the mitotype pool. Although there is a structure observed between infection status and mtDNA haplotype, the presence of two uninfected individuals in the main infected haplotype in *C. imicola* from South Africa is consistent with a symbiont showing imperfect vertical symbiont transmission, generating uninfected individuals from lineages with the dominant *Cardinium* infected mitotype.

**Figure 1.**
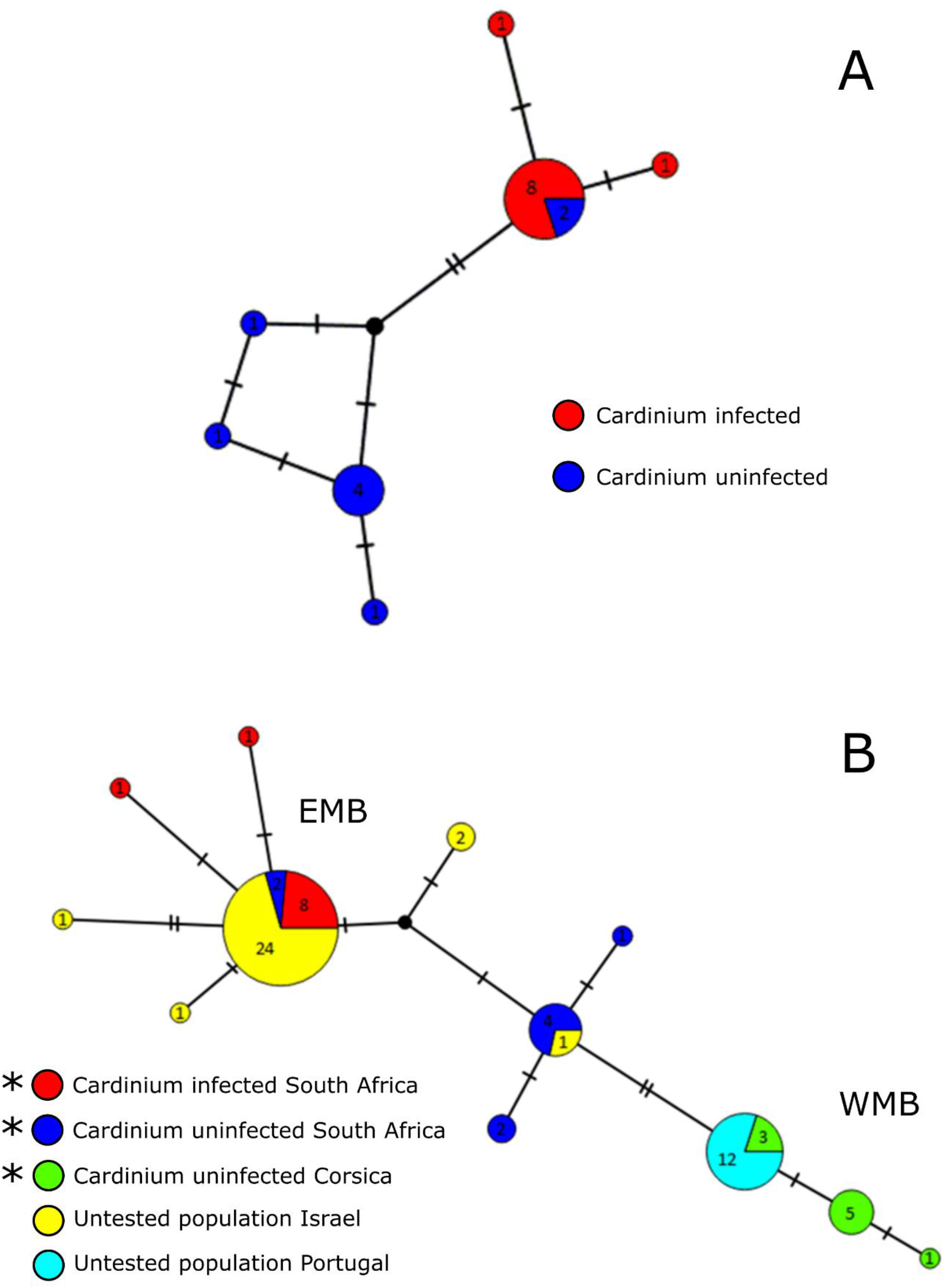
**A)** An mtDNA haplotype network of *Cardinium* infected and uninfected *Culicoides imicola* from a single site at ARC-OVI, Onderstepoort, South Africa; based on a 517 bp *COI* sequence. **B)** An mtDNA haplotype network of *Culicoides imicola* from sites spanning South Africa and the Mediterranean basin; based on a 451 bp *COI* sequence. Haplotype networks were generated using the TCS network algorithm in PopART v1.7 (Leigh & Bryant 2015). Numbers within circles represent the numbers of individuals designated to each haplotype. The numbers of substitutions separating haplotypes are indicated by dashes. EMB=Eastern Mediterranean Basin, WMB=Western Mediterranean Basin *Indicates mtDNA haplotypes generated in this study.

Extending the haplotype network to include *C. imicola* populations from the WMB (Corsica and Portugal) and the EMB (Israel), led to 12 haplotypes being observed with 0-1.77% range of divergence over a 451 bp region. Further to this, 24/29 Israeli (EMB) haplotypes clustered within the main *Cardinium* infected mitotype observed in South Africa, whereas the Portuguese (WMB) haplotypes all clustered within an uninfected Corsican mitotype. Although the infection status of these Portuguese and Israeli individuals have not been directly assessed, the recent Pagès (2017) screening for *Cardinium* found no indication of *C. imicola* infection in the Iberian Peninsula but others (5) have observed the symbiont to be common in Israel. In light of this, mitotype variation between the EMB and WMB mirrors infection heterogeneity between *C. imicola* populations from the EMB and WMB. Thus, mtDNA structure may be explained by the dispersal of *Cardinium* infected individuals exclusively into the EMB.

The matrilineal subdivision between *Cardinium* infected and uninfected *C. imicola* (Figure 1B) is similar to observations by previous studies (3,10) which noted a remarkable divergence between *C. imicola* mitotypes from the EMB and WMB. This observation suggests that the linkage disequilibrium of *Cardinium* and mitochondria reflects symbiont gene flow within the Mediterranean basin but may not assist in elucidating host gene flow. For example, if dispersal events occur involving *Cardinium*-infected *C. imicola*, selective sweeps of the symbiont may erase any previous biodiversity in this marker. This is reminiscent of similar patterns observed in other insects with endosymbiont infections: the mosquito vector, *Culex pipiens* (11), the parasitoid wasp, *Nasonia vitripennis* (12) and the ladybird, *Adalia bipunctata* (13). In all these cases, the frequency of mtDNA haplotypes are more closely associated with the endosymbionts in a population rather than geography.

The haplotype networks produced in this study (Figure 1) suggest *Cardinium* infection in *Culicoides* is another example where the presence of a symbiont impacts mtDNA based inference of population history. Previous work had indicated Israeli and Southern Africa samples grouped together on analysis of mtDNA variation but were distinct on analysis of microsatellite variation (1). We conclude that association of *Cardinium* and mtDNA provides a likely explanation for this discordance, and these data support the general importance of using multiple loci in phylogeographic analysis.

