## Supplemental Table S1 for "*Cardinium* symbiosis as a potential confounder of mtDNA based phylogeographic inference in *Culicoides imicola* (Diptera: Ceratopogonidae), a vector of veterinary viruses"

| **Target** | **Primer name** | **Sequence (5’-3’)** | **Tm (°C)** | **Size (bp)** | **Reference** |
| --- | --- | --- | --- | --- | --- |
| Cytochrome oxidase subunit 1 (*COI*) | LCO2198 | GGTCAACAAATCATAAAGATATTGG | 51 | 708 | Folmer *et al.* 1994 |
|  | HCO1490 | TAAACTTCAGGGTGACCAAAAAATCA | 56 |  |  |
| Gyrase B | gyrB23F | GGAGGATTACATGGYGTGGG | 60 | 1368 | Lewis *et al.* 2014 |
|  | gyrB1435R | GTAACGCTGTACATACACGGCATC | 60 |  |  |
|  | gyrBnest212F | AAGGCAACCCTATGCACCAA | 59 | 347 | This study |
|  | gyrBnest654R | GGYCTTAGTTTGCCCTTCAAATTG | 59 |  |  |

**Table S1.1.** *COI* and *Gyrase B* gene primer attributes.

| **Target** | **Initialisation** | **Denaturation, annealing, extension** | **Final extension, hold** | **Number of cycles** |
| --- | --- | --- | --- | --- |
| Cytochrome oxidase subunit 1 (COI) | 95°C/5min | 95°C/30s, 50°C/1min, 72°C/1min | 72°C/7min, 15°C/∞ | 35 |
| Gyrase B (Conventional) | 95°C/5min | 95°C/30s, 55°C/1min, 72°C/1min30s | 72°C/7min, 15°C/∞ | 35 |
| Gyrase B (Nested) | 95°C/5min | 95°C/25s, 55°C/30s, 72°C/45s | 72°C/7min, 15°C/∞ | 35 |

**Table S1.2.** PCR cycling conditions.
